# *De novo* genome assembly of the floating-hook moss *Warnstorfia fluitans*

**DOI:** 10.64898/2026.11.03.763626

**Authors:** Minh-Phuong Nguyen, Juan A. Galarza, Janne J. Koskimäki, Abrar Hussain Mian, Olga Podolich, Saija H.K. Ahonen, Anna Maria Pirttilä, Kaisa Lehosmaa

**Affiliations:** University of Oulu, Ecology and Genetics Research Unit, P.O. Box 3000, FI-90014, Oulu, Finland

## Abstract

The floating-hook moss *Warnstorfia fluitans* is a natural species dominant in subarctic and Arctic aquatic ecosystems. It is adapted to cold, acidic, and nutrient-deficient environments and possesses great potential for bioremediation by removing contaminants from polluted water in cold climate conditions. Here, we present the first *de novo* genome assembly of *W. fluitans* generated from pooled male and female gametophytes. The assembly is 725.04 Mb, has a N50 of 193.5 kb, a BUSCO completeness of 97.6% against the Viridiplantae lineage and 82% against the Embryophyta lineage. The whole assembly has Merqury quality value of 37 and Merqury completeness of 92%. A total of 30421 protein coding genes were predicted, with 78.62% completeness by compleasm and 87% OMArk completeness against the Embryophyta lineage. Repetitive sequences comprise 56.16% of the genome. We also present here the two complete circular chloroplast genomes and a linear mitochondrial genome of *W. fluitans*. This genomic resource will facilitate studies of ecological adaptation and a wide range of biological and evolutionary studies in bryophytes.

## Background & Summary

Aquatic bryophytes, such as mosses, are valuable phytoremediation agents to remove inorganic or organic pollutants from water due to their large surface area, ability to uptake elements through cell walls, and adaptation to harsh environments ^1–4^. The floating-hook moss *Warnstorfia fluitans* (Fig. 1), a dominant submerged moss found in subarctic and Arctic freshwater systems, is a promising candidate for remediation in nutrient-deficient, cold, and acidic water bodies where low temperature slows down the biodegradation process of pollutants ^5–7^. Previous studies have shown that *W. fluitans* can remove nitrogen and accumulate arsenic and iron from contaminated waters ^4,5,7,8^. These studies point out that the potential of *W. fluitans* for water remediation, especially in association with its microbial symbionts ^5^, is promising and can be further explored to treat polluted water systems in subarctic and Arctic environments. As the research on bryophyte genomics, evolution, ecology, and bioremediation-related traits is rapidly growing, a high-quality genome assembly of *W. fluitans* is needed. However, there is no genome assembly of the genus *Warnstorfia* or even closely related species or genera available to date.

**Fig. 1.**
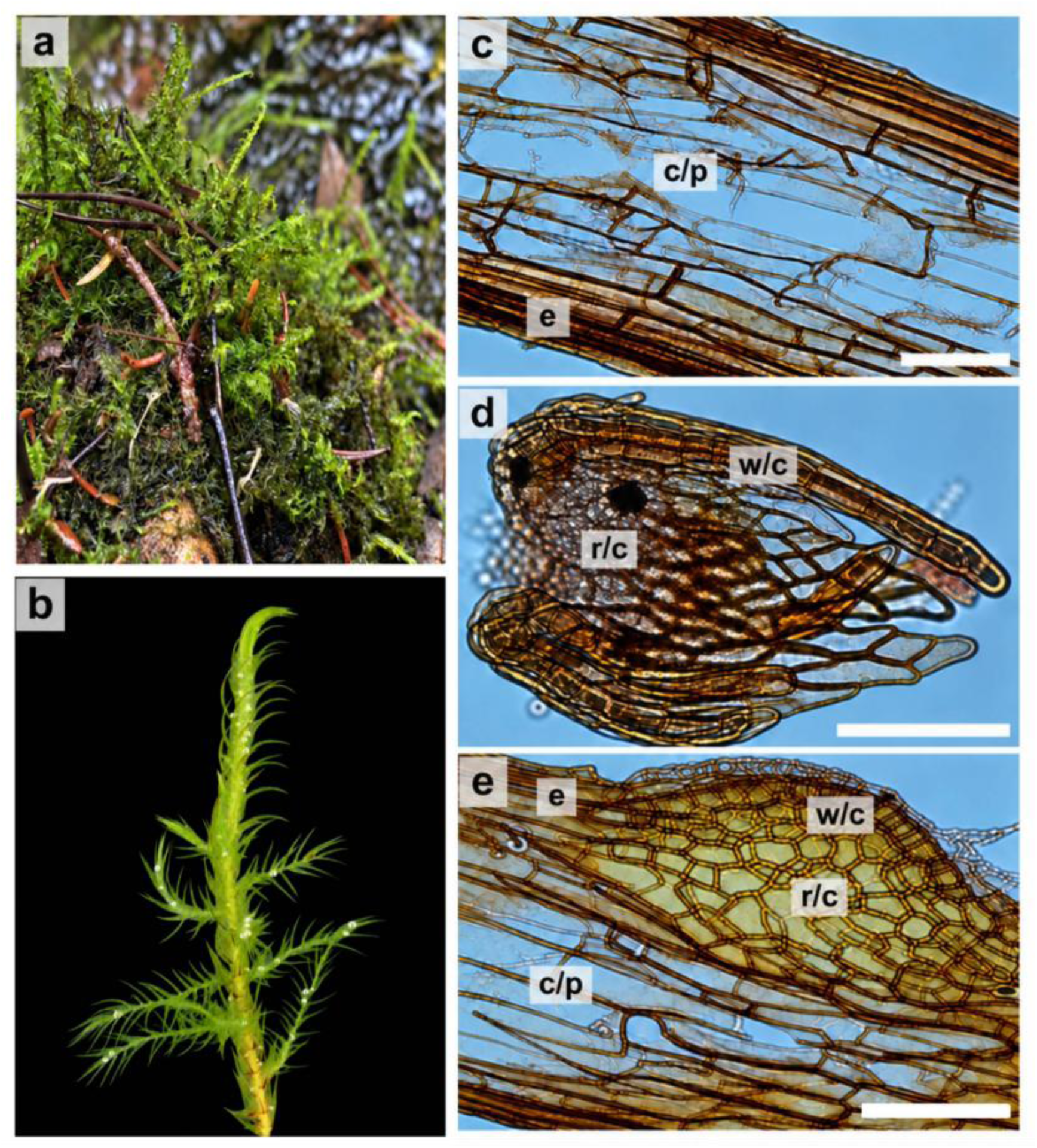
Morphological and microscopic features of the floating-hook moss *Warnstorfia fluitans*. (a) *W. fluitans* in its natural habitat. (b) A shoot of *W. fluitans*. (c) Microscopic view of the vegetative part. (d) Female reproductive structure (archegonium). (e) Male reproductive structure (antheridium). *c/p*, cortical parenchyma cells; *e*, epidermal cells; *w/c*, outer wall cells; *r/c*, inner reproductive cells. Scale bars: c–e = 50 µm.

We present here the first *de novo* genome assembly of *Warnstorfia* genus, focusing on the species *W. fluitans*. The final assembly was constructed from PacBio HiFi long reads and Illumina short reads originated from pooled male and female gametophytes and was annotated based on both transcript and protein evidence. The assembly spans 725.04 Mb with the N50 of 193.5 kb, and BUSCO completeness score of 97.6% against the Viridiplantae lineage and 82% against the Embryophyta lineage. Repetitive sequences accounted for 56.16% of the genome. We predicted 30421 protein coding genes using both transcriptomic and protein evidence. Our data provides a valuable genomic resource for a wide range of biological and evolutionary studies in bryophytes. It can serve the future studies in exploring gene functions and identifying candidate genes associated with important traits in this and closely related moss species for various practical applications including environmental remediation. The accompanying chloroplast and mitochondrial genomes from this study will be useful for phylogenetic comparative analysis among bryophytes.

## Methods

### Sample collection

We collected the moss *W. fluitans* from a manmade ditch in Pilpajärvi, Finland (64°58′10″ N, 25°48′03″ E) in autumn 2024 using a sterile rake. The moss material was grown in tap water in a flow-through bioreactor as described previously ^4^. The growth conditions consisted of a 16:8 h of light - dark cycle with a light intensity of 70 µmol m^-2^ s^-1^, a relative humidity of 70% and a temperature of 5°C. The inflow water rate was maintained at 1.2 mL min^-1^ using a peristaltic pump (MINIPULS 3, Gilson, Inc., Middleton, WI, USA).

### DNA extraction and whole genome sequencing

Twenty grams of whole moss plants, including both male and female gametophytes, were ground into a fine powder on a liquid nitrogen bath using a sterile mortar and pestle. The powder was divided into 100 eppendorfs, each containing 200 mg of the powder, for DNA extraction. The moss DNA was extracted from each tube following the protocol of Pirttilä et al. ^9^ with modification. Specifically, 900 µl of pre-heated (65°C) extraction buffer (2% CTAB, 2% PVP Mr 750,000, 2% β-mercaptoethanol, 2 M LiCl) was added to 200 mg of moss homogenized powder and incubated at 65°C for 20 min with mixing three times in between. Then the sample was extracted twice with one equal volume of chloroform:IAA (24:1, v:v). The DNA was precipitated in one equal volume of cold isopropanol and incubated for 1 h at room temperature. The DNA pellets were collected by centrifugation and washed three times with 500 µl 70% ethanol. DNA from all the extracted tubes was pooled into 1 mL of Tris-EDTA buffer (pH 8). RNA was digested from the pooled DNA sample by incubating the sample with 100 µg of PureLink RNase A (20 mg mL^-1^ Invitrogen, MA, USA) at 37 °C for 30 min. The DNA was then purified and sequenced at the BMKGENE sequencing company (Germany). A standard PacBio HiFi library preparation kit (SMRTbell Prep Kit 3.0) aiming at 10–20 kb circular consensus sequence with the desired insert size of approximately 15 kb was used. The library was sequenced on a single SMRT Cell on the PacBio Revio platform using a 30-h movie.

The same genomic DNA extracted above was purified using the SSTE buffer (1M NaCl, 0.5% SDS, 10 mM Tris-Cl, 1 mM EDTA, pH 8) and chloroform:IAA (24:1, v:v) following the protocol of Jaakola et al.^10^. The DNA was then purified one more time with AMPure XP magnetic beads (Beckman Coulter Life Sciences, IN, USA) following the manufacture’s instruction to produce high quantity and purity DNA for genomic sequencing with Illumina NovaSeq X Plus Series (PE150) at the Novogene sequencing company (Germany).

### Transcriptome sequencing

RNA samples were extracted from the moss material growing in tap water following the protocol of Jaakola et al. ^10^ with modification. Specifically, after precipitation of the RNA pellets using Lithium chloride (LiCl), DNA was digested from the RNA samples using DNase I (Fermentas, IN, USA) following the manufacturer’s instructions. Then RNA samples were purified following the rest of the protocol by Jaakola et al. ^10^. High purity and integrity RNA samples were sent for eukaryotic mRNA sequencing with Illumina NovaSeq X Plus Series (PE150) at Novogene sequencing company (Germany). A total of 44 Gb raw data was obtained for three transcriptomic replicates.

### Genome assembly

A total of 30 Gb of HiFi long-read and 50 Gb of Illumina short-read data were generated for the moss genome sequencing. We analyzed the quality of the HiFi long reads using Nanoplot v1.44.1 ^11^, removed the adapters using HifiAdapterFilt v3.0.0 ^12^, and filtered low quality and reads shorter than 1000 bp using cutadapt v4.9 ^13^. To remove contaminant reads from downstream analyses, we used Blobtools v1.1 ^14^, which employs both BLAST hits, coverage and GC content to classify the assembled contigs. Since our data originated from pooled male and female haploid genomes, we assembled the long reads using hifiasm v0.25.0 ^15^ with option −l0 to disable duplication purging. We used BLASTn megablast of BLAST+ v2.17.0 ^16^ against the nucleotide (nt) database and DIAMOND blastx v2.1.16 ^17^ against the Uniprot Reference Proteomes 2025.03 database to get the BLAST hits. We then employed minimap2 v2.28 ^18^ and SAMtools v1.21 ^19^ with the long reads to generate the read coverage of the initial assembled contigs. We filtered out the contaminant contigs with Blobtools for two rounds with the ‘bestsumorder’ rule and kept only the contigs that matched to “streptophyta” or had no matches (“no-hit”). We assembled the plastid and mitochondrial genomes from the cleaned long reads using Oatk ^20^ and separated the organelle reads from the nuclear long reads using SAMtools. We then built the moss nuclear genome using the nuclear long reads and hifiasm without duplication purging. Since our moss genome data was derived from both male and female haploids, there was no further duplication purging performed. We used Racon v1.5.0 ^21^ and the nuclear long reads for five rounds to correct the assembled genome.

For the Illumina short reads, we analyzed the quality using FastQC v0.12.1 ^22^, removed adapters, and trimmed reads based on quality using fastp v0.24.0 ^23^. Next, we aligned the short reads to the organelle long reads and nuclear long reads using BWA-MEM v0.7.18 ^24^ and extracted the organelle short reads and nuclear short reads using SAMtools. We used the nuclear short reads to correct the Racon-polished assembly with Pilon v1.24 ^25^ for five rounds. The final polished assembly was scaffolded using LRScaf v1.1.12 ^26^. The quality of the assembly was assessed by QUAST v5.3.0 ^27^, Merqury v1.4.1 ^28^, BUSCO v6.1.0 ^29^, and compleasm v0.2.9 ^30^ against the lineages Viridiplantae_odb12 and Embryophyta_odb12 after each step. Out of 1,379,545 long reads (30 Gb), 402,306 reads (9.6 Gb) remained after filtering out the contaminating reads with Blobtools, and among these reads, 4648 reads (1.14%) belonged to organelle genomes. 397,127 nuclear reads (9.48 Gb) were retained after further screening with the Foreign Contamination Screen (FCS) tool suite of the National Center for Biotechnology Information (NCBI) during the genome assembly submission to NCBI. The LRScaf-scaffolded assembly did not significantly improve the total BUSCO scores but increased the number of duplicated BUSCO orthologs compared to the Pilon-polished assembly (Supplementary Material, Table S1-S3). Therefore, the Pilon-polished genome assembly was chosen as the final one. The nuclear genome assembly had a mean coverage of 5.7x by the HiFi long reads for structural and repeat resolution and 15.6x by the Illumina reads for base-level correction and validation. The final assembly spanned 725.04 Mb, arranging in 5851 contigs with the contig N50 of 193.5 kb, had Merqury completeness of 91.92% and quality value (QV) of 37 (Table 1). Both BUSCOs and compleasm analyses showed the completeness of 97.6 % against the lineage Viridiplantae_odb12 and 82% for the lineage Embryophyta_odb12 (Table 2). The duplicated BUSCOs were high based on compleasm results, e.g., 83.45% for Viridiplantae and 66.29% for Embryophyta.

**Table 1:** The statistics of contig-level genome assembly of *Warnstorfia fluitans* based on QUAST and Merqury.

| **Genome statistic** | **Final assembly** |
|---|---|
| Total length | 725045309 |
| # of sequences | 5851 |
| # sequences (≥ 0 bp) | 5851 |
| # sequences (≥ 1000 bp) | 5851 |
| Total length (≥ 0 bp) | 725045309 |
| Total length (≥ 1000 bp) | 725045309 |
| Largest sequence | 1039778 |
| GC level | 37.77 |
| N50 | 193474 |
| N90 | 53141 |
| auN | 242337.9 |
| L50 | 1109 |
| L90 | 3932 |
| # N’s per 100 kbp | 0 |
| Completeness | 91.92 % |
| Quality value (QV) | 37 |

**Table 2:** Completeness assessment of the genome assembly *Warnstorfia fluitans* by BUSCO orthologs using compleasm and BUSCO.

|  | Viridiplantae_odb12 |  | Embryophyta_odb12 |  |
|---|---|---|---|---|
| Software | Compleasm | BUSCO | Compleasm | BUSCO |
| Complete buscos (C = S + D) | 802 (97.56%) | 802 (97.6%) | 1661 (81.98%) | 1665 (82.2%) |
| Complete and single copy buscos (S) | 116 (14.11%) | 188 (22.9%) | 318 (15.70%) | 383 (18.9%) |
| Complete and duplicated buscos (D) | 686 (83.45%) | 614 (74.7%) | 1343 (66.29%) | 1282 (63.3%) |
| Fragmented buscos, Subclass 1 (F) | 3 (0.36%) | 3 (0.4%) | 90 (4.44%) | 77 (3.8%) |
| Fragmented buscos, Subclass 2 (I) | 0 (0%) | 0 (0%) | 0 (0%) | 0 (0%) |
| Missing buscos (M) | 17 (2.07%) | 17 (2.1%) | 275 (13.57%) | 284 (14%) |

### Repeat annotation

We built the custom repeat elements with our final assembly using RepeatModeler v2.0.7 ^31^ with the long terminal repeat (LTR) discovery pipeline and soft-masked the assembly by using RepeatMasker v4.1.9 ^32^ and the *de novo* repeat library generated by RepeatModeler (with options --nolow to not hard-mask low complexity DNA or simple repeats, and -xsmall to soft-mask). Our analysis identified 407.22 Mb of repetitive sequences, representing 56.16% of the genome (Table 3). The classified portion is dominated by DNA transposons (6.94%) and long terminal repeat (LTR) elements (2.7%), particularly Ty1/Copia and Gypsy/DIRS1. 45.79% of the repeats was unclassified.

**Table 3:** The statistics of repeat sequences in *Warnstorfia fluitans* genome. The repeat sequences were identified and soft-masked using RepeatModeler and RepeatMasker.

| Sequences | 5851 |  |  |
|---|---|---|---|
| Total length | 725045309 bp |  |  |
| GC level | 37.77 % |  |  |
| Bases masked | 407221349 bp | (56.16 %) |  |
|  | Number of elements | length occupied (bp) | % of sequence |
| Retroelements | 21679 | 21559655 | 2.97 |
| LINEs: | 7231 | 2017064 | 0.28 |
| L1/CIN4 | 117 | 140594 | 0.02 |
| LTR elements: | 14448 | 19542591 | 2.70 |
| Ty1/Copia | 3360 | 5436560 | 0.75 |
| Gypsy/DIRS1 | 8407 | 13619577 | 1.88 |
| DNA transposons | 86689 | 50336026 | 6.94 |
| hobo-Activator | 8602 | 12497081 | 1.72 |
| Tc1-IS630-Pogo | 19949 | 5736036 | 0.79 |
| MULE-MuDR | 1315 | 1336450 | 0.18 |
| PiggyBac | 6199 | 2791108 | 0.38 |
| Tourist/Harbinger | 8806 | 2595651 | 0.36 |
| Rolling-circles | 3829 | 2947446 | 0.41 |
| Unclassified: | 920491 | 331966305 | 45.79 |
| Total interspersed repeats: |  | 403861986 | 55.70 |
| Small RNA: | 747 | 411917 | 0.06 |

### Protein-coding gene prediction and functional annotation

For the RNA-seq data, we analyzed the quality of the raw data using FastQC, performed kmer-based read corrections using rCorrector v1.0.7 ^33^, and removed unfixable read pairs with the python script *FilterUncorrectabledPEfastq*.py (https://github.com/harvardinformatics/TranscriptomeAssemblyTools/). We performed adaptor and poly-G trimming, and a light quality trimming from the data using fastp v0.24.0 ^23^ (with options: cut_window_size 4, cut_mean_quality 5). Reads originating from the rRNAs were removed with SortMeRNA v4.3.7 ^34^ and the SILVA database 138.2 (LSURef and SSURef NR99) ^35^. The filtered RNA-seq data was aligned to the soft-masked assembly using STAR v2.17.11b ^36^. We predicted genes in the masked assembly based on the alignment data generated by STAR and the protein database OrthoDB 12 partition Viridiplantae (available at https://bioinf.uni-greifswald.de/bioinf/partitioned_odb12/) using BRAKER3 ^37^. The completeness of the gene annotation was assessed by compleasm and BUSCO against both lineage datasets, Viridiplantae_odb12 and Embryophyta_odb12, utilizing the protein output file generated by BRAKER3. We also used OMArk v0.5.0 ^38^ and its constructed database LUCA.h5 to assess completeness, consistency, and contamination of the output file ^38^. Compleasm-based statistics showed that 95.01% and 78.62% of conserved orthologs are complete for both lineages (Table 4). OMArk analysis showed the overall completeness of 86.96% against the Embryophyta clade, and 86.38% of proteins showed consistent lineage placement (Table 5). The *W. fluitans* proteins were found consistent by comparison with the moss *Physcomitrium patens*, and no contaminant was detected by OMArk.

**Table 4:** BUSCO completeness assessment of the *Warnstorfia fluitans* predicted proteins.

|  | Viridiplantae_odb12 |  | Embryophyta_odb12 |  |
|---|---|---|---|---|
|  | compleasm | BUSCO | compleasm | BUSCO |
| Complete BUSCOs (C) | 781 (95.01%) | 781 (95%) | 1593 (78.62%) | 1590 (78.5%) |
| Complete and single-copy BUSCOs (S) | 102 (12.41%) | 112 (13.6%) | 259 (12.78%) | 259 (12.8%) |
| Complete and duplicated BUSCOs (D) | 679 (82.60%) | 669 (81.4%) | 1334 (65.84%) | 1331 (65.7%) |
| Fragmented BUSCOs (F) | 10 (1.22%) | 10 (1.2%) | 114 (5.63%) | 106 (5.2%) |
| Missing BUSCOs (M) | 31 (3.77%) | 31 (3.8%) | 319 (15.75%) | 330 (16.3%) |

**Table 5:** Summary of OMArk proteome quality assessment for *Warnstorfia fluitans* genome.

| Item | Count | % |
|---|---|---|
| Completeness assessment |  |  |
| Ancestral clade used | Embryophyta |  |
| Number of conserved HOGs | 6167 |  |
| Complete | 5364 | 86.98 % |
| Single: | 1960 | 31.78 % |
| Duplicated: | 3404 | 55.20 % |
| Duplicated, Unexpected | 3195 | 51.81 % |
| Duplicated, Expected | 209 | 3.39 % |
| Missing: | 803 | 13.02 % |
| Consistency assessment |  |  |
| Number of proteins | 30421 |  |
| Total consistent | 26322 | 86.53 % |
| Consistent, partial hits | 4896 | 16.09 % |
| Consistent, fragmented | 717 | 2.36 % |
| Inconsistent lineage placements |  |  |
| Total inconsistent | 1277 | 4.20 % |
| Inconsistent, partial hits | 454 | 1.49 % |
| Inconsistent, fragmented | 158 | 0.52 % |
| Contaminants |  |  |
| Total contaminants | 0 | 0 % |
| Contaminants, partial hits | 0 | 0 % |
| Contaminants, fragmented | 0 | 0 % |
| Total Unknown | 2822 | 9.28 % |
| Species composition |  |  |
| Clade | *Physcomitrium patens* |  |
| No. associated query proteins | 27599 | 90.72 % |

We assigned functions to the predicted proteins generated by BRAKER3 using the eggNOG-mapper v3.0.0-beta6 ^39–43^ and InterProScan v5.78-109.0 ^44^. In total, 93.47% of the predicted proteins were annotated in at least one of the databases by eggNOG-mapper (Table 6) and 90.62% were assigned InterPros assessions (Table 6).

**Table 6:** Overview of the annotated functions for *Warnstorfia fluitans* genome.

|  | Gene function annotation | no. annotated protein | % annotated proteins |
|---|---|---|---|
| eggNOG-mapper | Total annotated | 28436 | 93.47 |
|  | Clusters of Orthologous genes (COGs) | 9389 | 30.86 |
|  | Gene Ontology (GO) | 22685 | 74.57 |
|  | Enzyme Commission (EC) | 11186 | 36.77 |
|  | Kyoto Encyclopedia of Genes and Genomics (KEGG) | 20394 | 67.04 |
|  | Pfam | 24996 | 82.17 |
| InterProScan | InterPros | 27567 | 90.62 |
|  | Signature | 29771 | 97.86 |

### Organelle genomes assembly and annotation

We assembled both plastid and mitochondrial genomes from the decontaminated HiFi reads using Oatk with default settings and the pre-built organelle gene profile database of *embryophyte* version 20230921. The final organelle genomes were annotated and visualized with Chlorobox online software tools (https://chlorobox.mpimp-golm.mpg.de/), GeSeq ^45^ and OGDraw ^46^. For GeSeq, a circular sequence source was selected for the plastids, keeping only the best annotation. Annotation via ‘BLAT search’ (default settings) was done with GeSeq default MPI-MP reference set and NCBI RefSeq references belonging to *Calliergonella cuspidata* (NC_049070.1) and the well-annotated *Physcomitrium patens* (NC_005087.1). Among the available references in Geseq we chose *C. cuspidata* because it is the closest species to the family *Calliergonaceae*, to which the genus *Warnstorfia* belongs based on the phylogenetic tree produced previously ^47^. The BLAT search parameters included 85% protein search identity, as mentioned previously ^48^, and 85% rRNA, tRNA, DNA search identity. HMMER profile search, ARAGORN, and tRNAscan-SE were selected with default settings, as recommended. Since there was no reference mitochondrial genomes of the family *Calliergonaceae* available in the Geseq database, we selected members of the Hypnales order as references for the BLAT search, including *Sanionia uncinata* (NC_027974.1), *Brachythecium rivulare* (NC_031212.1), *Callicladium imponens* (NC_024516.1), *Climacium americanum* (NC_024515.1), *Pseudanomodon attenuatus* (NC_021931.1), and *Ptychomnion cygnisetum* (NC_024514.1). The annotations were summarized into GenBank files. Oatk resulted in two circular chloroplast genomes with the lengths of 124,894 bp (pltd_1, 201.5x mean coverage) and 124,856 bp (pltd_2, 82.6x mean coverage). In addition, a 103,712 bp linear mitochondrion (mito, 71.9x mean coverage) was obtained. The two plastids differ by length at the large single-copy (LSC), small single-copy (SSC), and inverted-repeat (IR) regions (Supplementary Material, Table S4). The GC ratio of the two plastids is 28.60% and 28.59%, respectively, similar to that of *P. patens* (28.5%) ^49^. They both contain four duplicated ribosomal RNA (rRNA), 32 tRNA, and 82 protein-coding genes in the same order (Table 7, Fig. 2). The chloroplast genome structure and gene order are highly similar to other bryophytes^48,49^. The linear mitochondrial genome (40.89% GC) contains the three rRNA, 24 tRNA, and 40 protein-coding genes necessary for a functional mitochondrion (Table 8, Fig. 3).

**Fig. 1:**
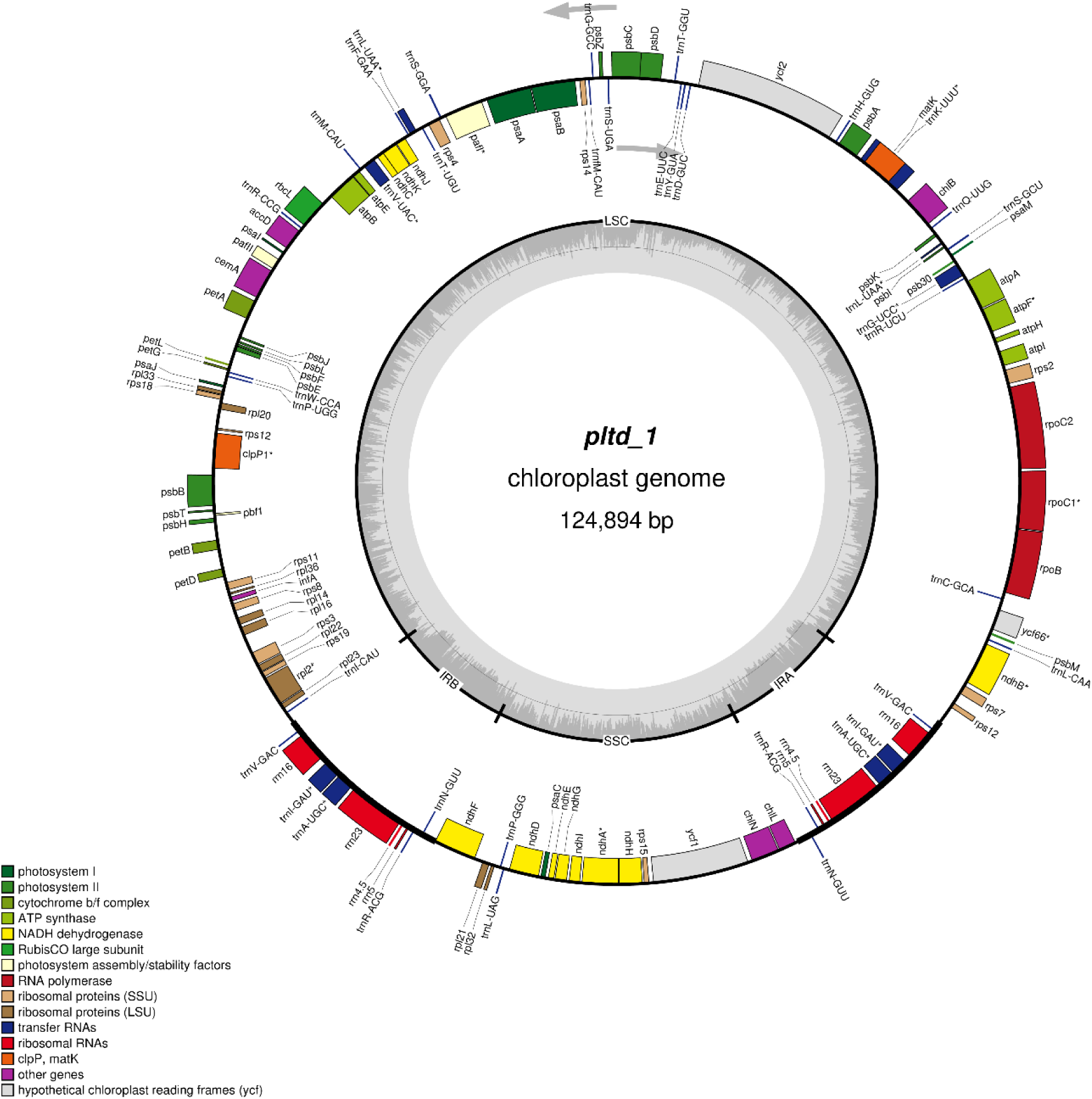
Circular map of the *Warnstorfia fluitans* chloroplast genome pltd_1. The large single-copy (LSC) and small single-copy (SSC) regions are separated by inverted repeats (IRA and IRB). Arrows indicate the transcription direction for genes outside (counterclockwise) and inside (clockwise) the map. Genes with related functions are shown in the same colour. Asterisks indicate genes having introns.

**Fig. 2:**
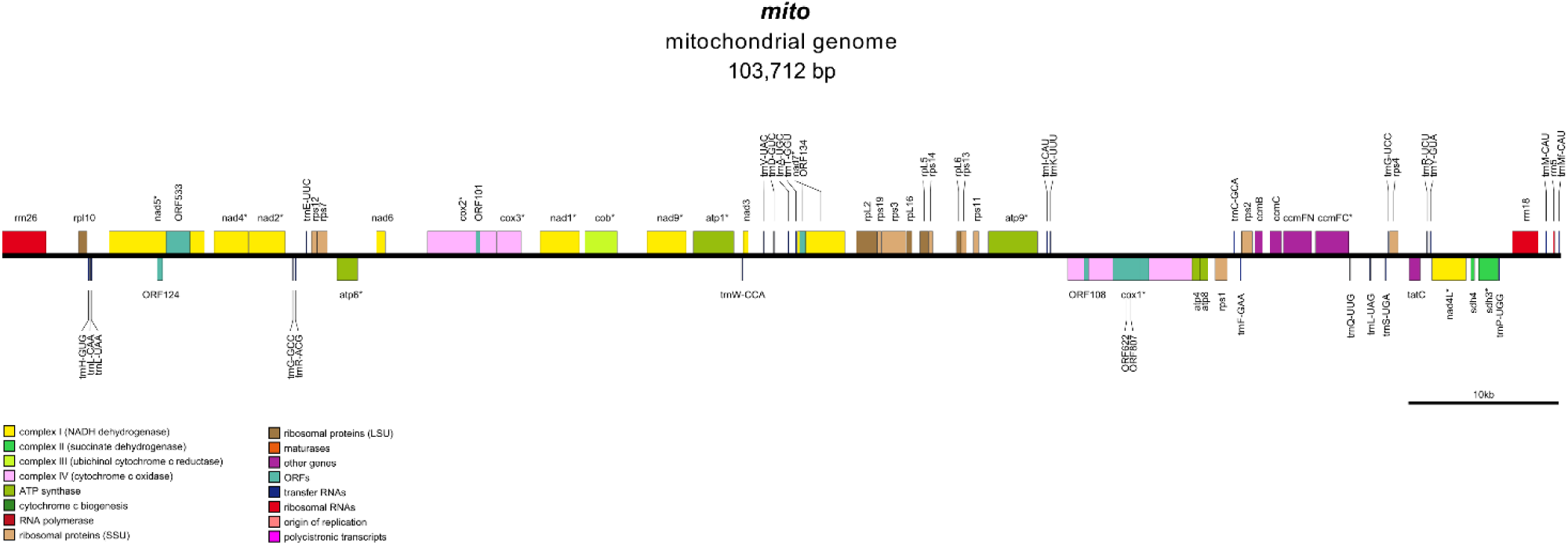
Map of the *Warnstorfia fluitans* mitochondrial genome.

**Table 7:** Genes in *Warnstorfia fluitans* plastid genomes.

| Gene category | Gene group | Gene name |
|---|---|---|
| Photosynthetic-related genes | ATP synthase | atpA, atpB, atpE, atpF<sup>a</sup>, atpH, atpI |
|  | Chlorophyll biosynthesis | chlB, chlL, chlN |
|  | Cytochrome b6/f | petA, petB, petD, petG, petL |
|  | NADH oxidoreductase | ndhA<sup>a</sup>, ndhB<sup>a</sup>, ndhC, ndhD, ndhE, ndhF<sup>c</sup>, ndhG, ndhH, ndhI, ndhJ, ndhK |
|  | Rubisco | rbcL |
|  | photosystem I | psaA, psaB, psaC, psaI, psaJ, psaM, pafI<sup>b</sup>, pafII |
|  | photosystem II | psbA, psbB, psbC, psbD, psbE, psbF, psbH, psbI, psbJ, psbK, psbL, psbM, psbT, psbZ, psb30, pbf1 |
| Self-replication genes (transcription and translation-related genes) | DNA-dependent RNA polymerase | rpoB, rpoC1<sup>a</sup>, rpoC2 |
|  | Large subunit ribosomal protein | rpl2<sup>a</sup>, rpl14, *rpl16*, rpl20, rpl21, rpl22, rpl23, rpl32, rpl33, rpl36 |
|  | Small subunit ribosomal protein | rps2, rps3, rps4, rps7, rps8, rps11, rps12<sup>d</sup>, rps14, rps15, rps18, rps19 |
|  | Ribosomal RNA genes | rrn4.5<sup>c</sup>, rrn5<sup>c</sup>, rrn16<sup>c</sup>, rrn23<sup>c</sup> |
|  | Transfer RNA genes | trnL-CAA, trnC-GCA, trnR-UCU, trnG-UCC<sup>a</sup>, trnS-GCU, trnL-UAA<sup>a</sup>, trnQ-UUG, trnK-UUU<sup>a</sup>, trnH-GUG, trnD-GUC, trnY-GUA, trnE-UUC, trnT-GGU, trnS-UGA, trnG-GCC, trnfM-CAU, trnS-GGA, trnT-UGU, trnF-GAA, trnV-UAC<sup>a</sup>, trnM-CAU, trnR-CCG, trnW-CCA, trnP-UGG, trnI-CAU, trnV-GAC<sup>c</sup>, trnI-GAU<sup>ac</sup>, trnA-UGC<sup>ac</sup>, trnR-ACG<sup>c</sup>, trnN-GUU<sup>c</sup>, trnP-GGG, trnL-UAG |
| Other genes | Maturase | matK |
|  | Proteolysis | clpP1<sup>b</sup> |
|  | Carbon metabolism | cemA |
|  | Fatty acid synthesis | accD |
|  | Transfer initiation factor | infA |
| Proteins of unknown function | Conserved open reading frames | ycf1, 2, 66<sup>a</sup> |

<sup>a</sup>: gene contains 1 intron; <sup>b</sup>: genes contain two introns; <sup>c</sup>: Gene duplicated in IRs; <sup>d</sup>: trans-splicing gene

**Table 8:** Genes in *Warnstorfia fluitans* mitochondrial genome.

| Gene category | Gene group | Gene name |
|---|---|---|
| Oxidative phosphorylation / Respiratory chain | NADH dehydrogenase (complex I) | nad1<sup>b</sup>, nad2<sup>a</sup>, nad3, nad4<sup>a</sup>, nad4L<sup>a</sup>, nad5<sup>c</sup>, nad6, nad7<sup>b</sup>, nad9<sup>a</sup> |
|  | Succinate dehydrogenase (complex II) | sdh3<sup>a</sup>, sdh4 |
|  | Cytochrome b (complex III) | cob<sup>a</sup> |
|  | Cytochrome c oxidase (complex IV) | cox1<sup>c</sup>, cox2<sup>c</sup>, cox3<sup>a</sup> |
|  | ATP synthase (complex V) | atp1<sup>a</sup>, atp4, atp6<sup>a</sup>, atp8, atp9<sup>b</sup> |
|  | Cytochrome c biogenesis | ccmB, ccmC, ccmFC<sup>a</sup>, ccmFN |
| Mitochondrial gene expression and protein synthesis | Large subunit ribosomal protein (LSU) | rpl2, rpl5, rpl6, rpl10, rpl16 |
|  | Small subunit ribosomal protein (SSU) | rps1, rps2, rps3, rps4, rps7, rps11, rps12, rps13, rps14, rps19 |
|  | Ribosomal RNA genes | rrn5, rrn18, rrn26 |
|  | Transfer RNA genes | trnH-GUG, trnL-CAA, trnL-UAA, trnG-GCC, trnR-ACG, trnE-UUC, trnW-CCA, trnV-UAC, trnD-GUC, trnA-UGC, trnT-GGU, trnI-CAU, trnK-UUU, trnC-GCA, trnF-GAA, trnQ-UUG, trnL-UAG, trnS-UGA, trnG-UCC, trnR-UCU, trnY-GUA, trnP-UGG, trnM-CAU, trnMf-CAU |
| Mitochondrial protein transport | Protein translocation | tatC |
| Proteins of unknown function | Conserved ORFs | ORF101, ORF108, ORF124, ORF134, ORF533, ORF622, ORF807 |

<sup>a</sup>: gene contains 1 intron; <sup>b</sup>: gene contains two introns; <sup>c</sup>: gene contains 3 introns

### Data Records

The sequencing data from HiFi and Illumina were deposited to the NCBI SRA under the accession number PRJNA1469967. The nuclear and organelle genome assembly results are available in DDBJ/ENA/GenBank under the accession JCBAOS000000000. The version described in this paper is the version JCBAOS010000000. The soft-masked nuclear assembly and organelle assemblies, as well as their annotations can be accessed on Zenodo (https://doi.org/10.5281/zenodo.23725012).

### Technical Validation

During the two rounds of contamination screening with Blobtools, we used stringent parameters for both BLASTn (max_target_seqs 10, evalue 1e-25) and DIAMOND blastx (sensitive mode, evalue 1e-25) to classify the assembled contigs. The contigs were finally screened for contamination using the NCBI FCS tool suite. These screenings helped us retain only HiFi reads/contigs classified as Streptophyta or no-hit (Supplementary Material, Table S5). We used both Benchmarking Universal Single-copy Orthologs (BUSCO) and compleasm with both lineages Viridiplantae_odb12 and Embryophyta_odb12 to evaluate the completeness of the genome assembly. We compared the genome statistics and completeness at each polishing and scaffolding step in order to select the Pilon-polished genome as the final assembly (Supplementary Material, Table S1-S3). We employed Merqury to assess the accuracy of the final assembly, yielding the quality values (QV) of 37 and completeness rate of 91.92%. We checked the coverage across the assembly by aligning the nuclear Hifi reads to the final assembly. The coverage was relatively consistent across the assembly with a mean coverage of 5.73x (Supplementary Material, Fig. S1). We used MultiQC v1.35 ^50^, SAMtools, and BEDTools v2.31.1 ^51^ to summarize the alignment statistics when mapping the transcriptomic data to the genome assembly. An average of 85% of the RNA read pairs properly mapped to the assembly (Supplementary Material, Table S6), and 86% of the total contigs were covered by ≥50 read pairs, indicating that the transcriptomic data aligned consistently across the assembled genome. An average of 12.6% alignments (including all kinds of alignments: primary, secondary, or multimapping) mapped to the repeat regions of the soft-masked assembly (Supplementary Material, Table S6), indicating that the majority of the transcriptomic data aligned to non-repetitive regions. We used OMArk as the complementary method for proteomic assessment, which showed that the integrity of the predicted proteins was 86.98%, indicating a high level of genome completeness for *W. fluitans*. To examine the high duplicated rate of the predicted proteome, we extracted the longest isoforms using the script *print_longest_isoform.py* (https://github.com/gatech-genemark/ProtHint; ^52^ and analyzed the completeness of the longest isoforms using compleasm and OMArk. Both methods gave similar results for the longest isoforms compared to those for the whole set of predicted proteins (Supplementary Material, Table S7), indicating that the high duplication is likely not due to the presence of multiple isoforms. Based on the OMArk’s result, we selected 80 Hierarchical Orthologous Groups (HOGs), including 20 HOGs having the highest amount of duplicated genes, 40 HOGs having 3-6 duplicated genes, and 20 HOGs having two duplicated genes to further investigate the duplication rate. We extracted the DNA sequence of each duplicated gene including 50 kb of flanking regions on both ends of the gene (so called gene-flanking regions) using BEDTools. We then performed multiple sequence alignment for the gene-flanking regions of each HOG using the fast method FFT-NS-2 of MAFFT ^53^ to select the closest pairs of gene-flanking regions. A further pairwise alignment was done for the closest pairs using the function *nucmer* and visualized with dotplot using the function *mummerplot* of MUMmer4 ^54^. Among the 135 examined closest pairs, there were pairs showing disrupted diagonal or scattered alignment patterns, potentially reflecting genuine gene duplications (Fig. S2). On the other hand, a few pairs showed high collinearity and similarity, represented by diagonal dotplots, which may reflect either genomic duplications, or retaining of both male and female haploid genomes (Fig. S2). The HiFi coverage of these genes and gene-flanking regions were generally equal or higher than the mean coverage of the genome assembly (5.73x) (Table S8), suggesting that the majority of these gene-flanking regions are likely genomic duplications.

We examined the completeness of the Oatk-assembled organelle genomes by separating the HiFi reads for each and by aligning the reads back to their respective organelle genome using minimap2. Finally, we examined the alignments with the Intergrative Genomics Viewer (IGV) ^55^. The plastid HiFi reads provided a complete coverage of the plastid genomes for both pltd_1 and pltd_2, with the highest coverage values observed in the inverted-repeat regions. The mitochondrial HiFi reads were evenly distributed across the mitochondrial genome. We examined the Geseq-annotations for the organelle genomes using Artemis v18.2.0 ^56^, validated open reading frames (ORFs) by the ORF Finder of NCBI, and aligned the protein sequences using the NCBI BLASTp ^57^. The Geseq-annotated organelle genes were first analyzed for the start and stop codons using Artemis with the NCBI translation table 11 intended for bacterial, archaeal and plant plastid. In case the different annotation tools (BLAT, HMMER profile search, ARAGORN, and tRNAscan-SE) resulted in different coordinates for the same organelle genes, we prioritized the coordinates firstly identified by the BLAT or secondly shared by several tools. Among the BLAT hits, we prioritized the one with a higher coverage and matching identity. For the plastid genes, the ones that did not pass the validation by Artemis were subjected to ORF analysis using the NCBI ORF Finder, and the protein sequences of the longest ORFs for each gene were aligned against the corresponding proteins of the reference genome, *C. cuspidata*, using BLASTp. The validations confirmed that the structures and protein sequences of the ambiguous genes were highly similar to those of their corresponding reference genes, suggesting that these genes are likely complete and functional.

## Data Availability

The datasets generated and analyzed in this study are publicly available. The raw sequencing data are in NCBI archive under accession number PRJNA1469967. The assembled nuclear and organelle genome can be downloaded from NCBI using the accession number JCBAOS000000000. The version described in this paper is JCBAOS010000000. The soft-masked nuclear assembly and organelle assemblies, and their corresponding annotations are available on Zenodo (https://doi.org/10.5281/zenodo.23725012).

## Code Availability

All commands and pipelines used in the data analyses were performed according to the manuals provided by the respective tools used in this study. No specific or custom code was used.

## Author contributions

JK, AHM, OP, SHKA, AMP, and KL collected the moss material. AHM cultivated and prepared the moss material for DNA extraction. JJK assisted in DNA extraction and purification. OP performed morphological characterization based on microscopy. SHKA aided in sample collection and preparation of the moss materials for RNA sequencing. MPN performed DNA and RNA extraction, genome assembly and analyses. JAG provided guidance in the genome assembly pipeline and result interpretation. AMP and KL designed the study and acquired the funding. All authors contributed to the manuscript preparation.

## Competing Interests

The authors declare no competing interests.

## Acknowledgements

The authors thank Finnish Centre for Scientific Computing (CSC) for computational resources. We thank Stefan Prost and Zsófia Fekete for useful discussions for the genome analyses.

## Funding

This work was funded by the Research Council of Finland (RCF) under the MoPoW (24303864) and MOREWA (372737) projects. Also, the Kvantum Institute and Innovation Centre of the University of Oulu supported the work.

## Ethics statement

No ethical approval was required for this study as it did not involve human or animal subjects.

## Notes

### Competing Interest Statement

The authors have declared no competing interest.

