## Supplementary Material for "*De novo* genome assembly of the floating-hook moss *Warnstorfia fluitans*"

**Supplementary Figures:**

Fig. S1: Histogram of the contig coverage

Fig. S2: Dotplots of pairwise alignment for 135 pairs of duplicated gene-flanking regions (provided in a separate file). The pairs in red boxes are highly similar in pairwise alignment.

**Supplementary Tables:**

Table S1: Contig-level genome assembly statistics for Warnstorfia fluitans, assessed using QUAST, for the 5-round Racon, 5-round Pilon, LRScaf-scaffolded, and final version deposited in NCBI.

Table S2: BUSCO gene scores generated with compleasm and BUSCO based on 822 expected orthologs from the lineage Viridiplantae_odb12 for the 5-round Racon, 5-round Pilon, LRScaf-scaffolded, and final versions deposited in NCBI.

Table S3: BUSCO gene scores generated with compleasm and BUSCO based on 2026 expected orthologs of the lineage Embryophyte_odb12 for the 5-round Racon, 5-round Pilon, LRScaf-scaffolded, and final versions deposited in NCBI.

Table S4: Length of each region in the two plastid genomes of Warnstorfia fluitans.

Table S5: Taxonomic classification at two rounds of Blobtools using the “bestsumorder” rule and FCS assessment.

Table S6: The alignment statistics of the transcriptomic data mapped to the assembly generated by MultiQC, SAMtools and BEDTools.

Table S7: Summary of OMArk proteome quality assessment for longest isoforms of Warnstorfia fluitans

Table S8: Coverage of 135 pairs of duplicated gene-flanking regions (provided in a separate file). Coverage of highly similar pairs (i.e., those marked in red boxes in Fig. S2) is highlighted in purple.

**Supplementary Figures**

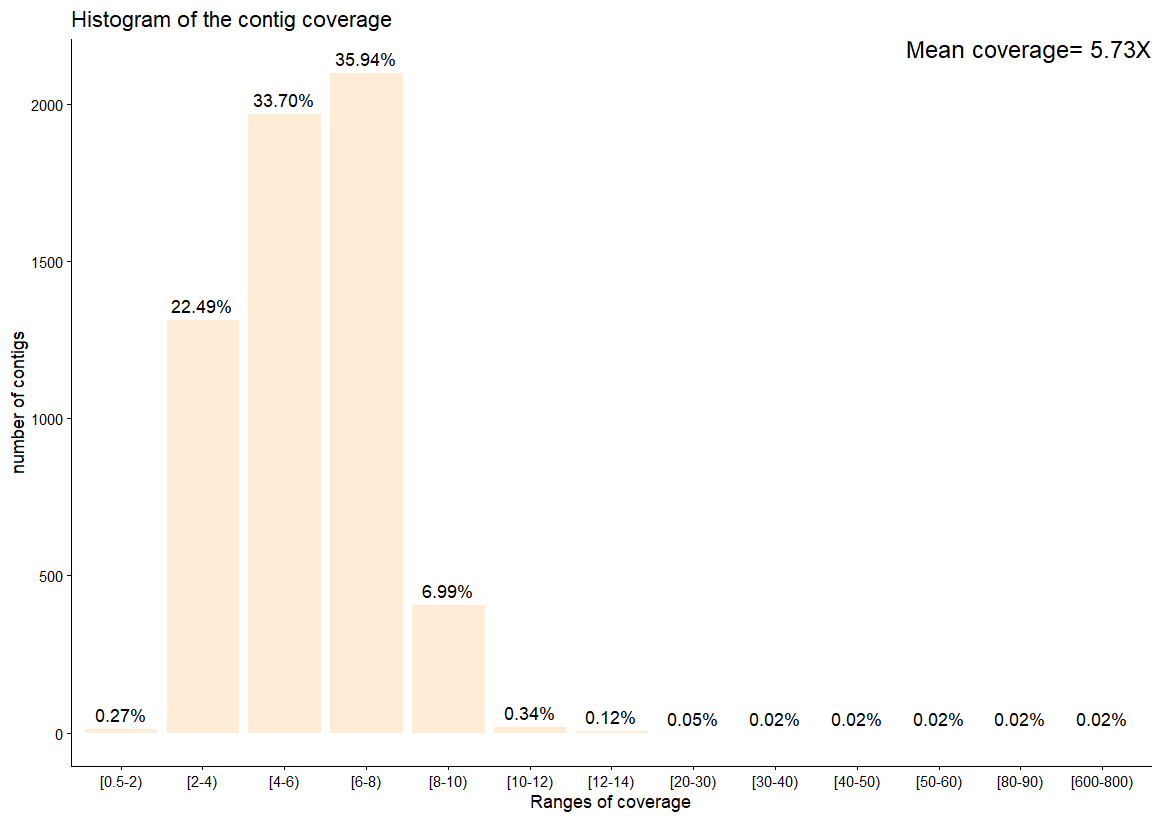

Fig. S1: Histogram of the contig coverage

Fig. S2: Dotplots of pairwise alignment for 135 pairs of duplicated gene-flanking regions (provided in a separate file). The pairs in red boxes are highly similar in pairwise alignment.

**Supplementary Tables**

Table S1: Contig-level genome assembly statistics for Warnstorfia fluitans, assessed using QUAST, for the 5-round Racon, 5-round Pilon, LRScaf-scaffolded, and final version deposited in NCBI.

| Genome statistic |  | Racon |  | Pilon |  | LRScaf |  | final |
| --- | --- | --- | --- | --- | --- | --- | --- | --- |
| Total length |  | 725834009 |  | 725613951 |  | 743997802 |  | 725045309 |
| # of sequences |  | 5855 |  | 5855 |  | 4221 |  | 5851 |
| # sequences (≥ 0 bp) |  | 5855 |  | 5855 |  | 4221 |  | 5851 |
| # sequences (≥ 1000 bp) |  | 5855 |  | 5855 |  | 4221 |  | 5851 |
| Total length (≥ 0 bp) |  | 725834009 |  | 725613951 |  | 743997802 |  | 725045309 |
| Total length (≥ 1000 bp) |  | 725834009 |  | 725613951 |  | 743997802 |  | 725045309 |
| Largest sequence |  | 1039783 |  | 1039778 |  | 1939788 |  | 1039778 |
| GC level |  | 37.78 |  | 37.78 |  | 37.78 |  | 37.77 |
| N50 |  | 193563 |  | 193556 |  | 341998 |  | 193474 |
| N90 |  | 53183 |  | 53168 |  | 68064 |  | 53141 |
| auN |  | 242424.4 |  | 242361.1 |  | 413904.7 |  | 242337.9 |
| L50 |  | 1109 |  | 1109 |  | 653 |  | 1109 |
| L90 |  | 3934 |  | 3934 |  | 2510 |  | 3932 |
| # N’s per 100 kbp |  | 0 |  | 0 |  | 517.53 |  | 0 |

Table S2: BUSCO gene scores generated with compleasm and BUSCO based on 822 expected orthologs from the lineage Viridiplantae_odb12 for the 5-round Racon, 5-round Pilon, LRScaf-scaffolded, and final versions deposited in NCBI.

| Software | Compleasm |  |  |  | BUSCO |  |  |  |
| --- | --- | --- | --- | --- | --- | --- | --- | --- |
| Assembly | Racon | Pilon | LRScaf | final | Racon | Pilon | LRScaf | final |
| Complete buscos (C = S + D) | 802 (97.56%) | 802 (97.56%) | 802 (97.56%) | 802 (97.56%) | 802 (97.6%) | 802 (97.6%) | 802 (97.6%) | 802 (97.6%) |
| Complete and single copy buscos (S) | 115 (13.99%) | 116 (14.11%) | 113 (13.75%) | 116 (14.11%) | 190 (23.1%) | 188 (22.9%) | 188 (22.9%) | 188 (22.9%) |
| Complete and duplicated buscos (D) | 687 (83.58%) | 686 (83.45%) | 689 (83.82%) | 686 (83.45%) | 612 (74.5%) | 614 (74.7%) | 614 (74.7%) | 614 (74.7%) |
| Fragmented buscos, Subclass 1 (F) | 3 (0.36%) | 3 (0.36%) | 3 (0.36%) | 3 (0.36%) | 3 (0.4%) | 3 (0.4%) | 3 (0.4%) | 3 (0.4%) |
| Fragmented buscos, Subclass 2 (I) | 0 (0%) | 0 (0%) | 0 (0%) | 0 (0%) | 0 (0%) | 0 (0%) | 0 (0%) | 0 (0%) |
| Missing buscos (M) | 17 (2.07%) | 17 (2.07%) | 17 (2.07%) | 17 (2.07%) | 17 (2.1%) | 17 (2.1%) | 17 (2.1%) | 17 (2.1%) |

Table S3: BUSCO gene scores generated with compleasm and BUSCO based on 2026 expected orthologs of the lineage Embryophyte_odb12 for the 5-round Racon, 5-round Pilon, LRScaf-scaffolded, and final versions deposited in NCBI.

| Software | Compleasm |  |  |  | BUSCO |  |  |  |
| --- | --- | --- | --- | --- | --- | --- | --- | --- |
| Assembly | Racon | Pilon | LRScaf | final | Racon | Pilon | LRScaf | final |
| Complete buscos (C = S + D) | 1658 (81.84%) | 1661 (81.98%) | 1662 (82.03%) | 1661 (81.98%) | 1663 (82.1%) | 1665 (82.2%) | 1664 (82.1%) | 1665 (82.2%) |
| Complete and single copy buscos (S) | 322 (15.89%) | 318 (15.70%) | 302 (14.91%) | 318 (15.70%) | 386 (19.1%) | 383 (18.9%) | 370 (18.3%) | 383 (18.9%) |
| Complete and duplicated buscos (D) | 1336 (65.94%) | 1343 (66.29%) | 1360 (67.13%) | 1343 (66.29%) | 1277 (63%) | 1282 (63.3%) | 1294 (63.9%) | 1282 (63.3%) |
| Fragmented buscos, Subclass 1 (F) | 92 (4.54%) | 90 (4.44%) | 89 (4.39%) | 90 (4.44%) | 77(3.8%) | 77 (3.8%) | 79 (3.9%) | 77 (3.8%) |
| Fragmented buscos, Subclass 2 (I) | 0 (0%) | 0 (0%) | 0 (0%) | 0 (0%) | 0 (0%) | 0 (0%) | 0 (0%) | 0 (0%) |
| Missing buscos (M) | 276 (13.62%) | 275 (13.57%) | 275 (13.75%) | 275 (13.57%) | 286 (14.1%) | 284 (14%) | 283 (14%) | 284 (14%) |

Table S4: Length of each region in the two plastid genomes of Warnstorfia fluitans.

|  | pltd_1 | pltd_2 |
| --- | --- | --- |
| Large single-copy (LSC) | 87854 bp | 87796 bp |
| Small single-copy (SSC) | 18498 bp | 18538 bp |
| Inverted-repeat region A (IRA) | 9271 bp | 9261 bp |
| Inverted-repeat region B (IRB) | 9271bp | 9261 bp |

Table S5: Taxonomic classification at two rounds of Blobtools using the “bestsumorder” rule and FCS assessment. Bold texts indicate the selected contigs (including chloroplasts and mitochondrion) assigned to Warnstorfia fluitans. Asterisk indicates four contigs classified as fungal, which were removed, and one contig with a region classified as fungal, which was trimmed in the final version deposited in NCBI.

| Blobtools round1 taxa | No. contigs |  | Blobtools round2 taxa | No. contigs |  | FCS assessment | No. contigs |
| --- | --- | --- | --- | --- | --- | --- | --- |
| **Streptophyta** | **5462** |  | **Streptophyta** | **5602** |  | **Streptophyta** | **5852** |
| Pseudomonadota | 2664 |  | **no-hit** | **179** |  | Fungi* | 5 |
| Bacteroidota | 1592 |  | Chlorophyta | 33 |  | Bacteria | 1 |
| Verrucomicrobiota | 873 |  | Ciliophora | 10 |  |  |  |
| Cyanobacteriota | 506 |  | cellular organisms-undef | 9 |  |  |  |
| **no-hit** | **335** |  | Pseudomonadota | 8 |  |  |  |
| Ascomycota | 247 |  | Basidiomycota | 4 |  |  |  |
| undef | 226 |  | Oomycota | 4 |  |  |  |
| Planctomycetota | 224 |  | Bacteroidota | 2 |  |  |  |
| Basidiomycota | 210 |  | Chloroflexota | 2 |  |  |  |
| Acidobacteriota | 207 |  | Cyanobacteriota | 2 |  |  |  |
| Actinomycetota | 164 |  | Echinodermata | 2 |  |  |  |
| Chlamydiota | 87 |  | Heterolobosea | 2 |  |  |  |
| Myxococcota | 85 |  | Mollusca | 2 |  |  |  |
| Chlorophyta | 39 |  | Nematoda | 2 |  |  |  |
| Armatimonadota | 37 |  | Verrucomicrobiota | 2 |  |  |  |
| Uroviricota | 35 |  | Actinomycetota | 1 |  |  |  |
| Chlorobiota | 32 |  | Ascomycota | 1 |  |  |  |
| Arthropoda | 31 |  | Bacillariophyta | 1 |  |  |  |
| Chloroflexota | 25 |  | Brachiopoda | 1 |  |  |  |
| Candidatus Melainabacteria | 23 |  | Cnidaria | 1 |  |  |  |
| Nucleocytoviricota | 21 |  | Discosea | 1 |  |  |  |
| Oomycota | 17 |  | Euglenozoa | 1 |  |  |  |
| Bacillota | 16 |  | Nucleocytoviricota | 1 |  |  |  |
| Chordata | 14 |  |  |  |  |  |  |
| Thermodesulfobacteriota | 13 |  |  |  |  |  |  |
| Candidatus Thorarchaeota | 12 |  |  |  |  |  |  |
| Mucoromycota | 11 |  |  |  |  |  |  |
| Rotifera | 11 |  |  |  |  |  |  |
| Ciliophora | 10 |  |  |  |  |  |  |
| Cnidaria | 8 |  |  |  |  |  |  |
| Candidatus Nomuraibacteriota | 7 |  |  |  |  |  |  |
| Gemmatimonadota | 6 |  |  |  |  |  |  |
| Bacillariophyta | 5 |  |  |  |  |  |  |
| Bdellovibrionota | 5 |  |  |  |  |  |  |
| Evosea | 5 |  |  |  |  |  |  |
| Nitrospirota | 5 |  |  |  |  |  |  |
| Mollusca | 4 |  |  |  |  |  |  |
| Nematoda | 4 |  |  |  |  |  |  |
| Apicomplexa | 3 |  |  |  |  |  |  |
| Candidatus Rokuibacteriota | 3 |  |  |  |  |  |  |
| Platyhelminthes | 3 |  |  |  |  |  |  |
| Rhodophyta | 3 |  |  |  |  |  |  |
| Thermoproteota | 3 |  |  |  |  |  |  |
| Campylobacterota | 2 |  |  |  |  |  |  |
| Ignavibacteriota | 2 |  |  |  |  |  |  |
| Lentisphaerota | 2 |  |  |  |  |  |  |
| Spirochaetota | 2 |  |  |  |  |  |  |
| Tardigrada | 2 |  |  |  |  |  |  |
| Zoopagomycota | 2 |  |  |  |  |  |  |
| Brachiopoda | 1 |  |  |  |  |  |  |
| Candidatus Bathyarchaeota | 1 |  |  |  |  |  |  |
| Candidatus Binatota | 1 |  |  |  |  |  |  |
| Candidatus Dormiibacterota | 1 |  |  |  |  |  |  |
| Candidatus Iainarchaeota | 1 |  |  |  |  |  |  |
| Candidatus Kaiseribacteriota | 1 |  |  |  |  |  |  |
| Candidatus Latescibacterota | 1 |  |  |  |  |  |  |
| Candidatus Moraniibacteriota | 1 |  |  |  |  |  |  |
| Candidatus Poribacteria | 1 |  |  |  |  |  |  |
| Candidatus Saccharimonadota | 1 |  |  |  |  |  |  |
| Candidatus Tayloriibacteriota | 1 |  |  |  |  |  |  |
| Candidatus Uhriibacteriota | 1 |  |  |  |  |  |  |
| Candidatus Wolfeibacteriota | 1 |  |  |  |  |  |  |
| Candidatus Yonathiibacteriota | 1 |  |  |  |  |  |  |
| Candidatus Zambryskiibacteriota | 1 |  |  |  |  |  |  |
| Chytridiomycota | 1 |  |  |  |  |  |  |
| Deinococcota | 1 |  |  |  |  |  |  |
| Discosea | 1 |  |  |  |  |  |  |
| Echinodermata | 1 |  |  |  |  |  |  |
| Euglenozoa | 1 |  |  |  |  |  |  |
| Haptophyta | 1 |  |  |  |  |  |  |
| Kiritimatiellota | 1 |  |  |  |  |  |  |
| Porifera | 1 |  |  |  |  |  |  |
| Preplasmiviricota | 1 |  |  |  |  |  |  |
| Rhodothermota | 1 |  |  |  |  |  |  |
| Tubulinea | 1 |  |  |  |  |  |  |
| candidate division WWE3 bacterium | 1 |  |  |  |  |  |  |

Table S6: The alignment statistics of the transcriptomic data mapped to the assembly generated by MultiQC, SAMtools and BEDTools. “Read” refers to forward and reverse RNA reads. “Alignments” includes all types of mappings (primary, secondary, and multimapping). Therefore, the number of alignments is greater than the number of reads.

|  | **Reads mapped to the assembly** | | | | | | | | | | **Alignments mapped to the repeat regions** | | |
| --- | --- | --- | --- | --- | --- | --- | --- | --- | --- | --- | --- | --- | --- |
| **RNA Sample** | **Total reads (million)** | **% aligned** | **% unique aligned** | **avg. Mapped length** | **annotated splices** | **mismatch rate** | **delete rate** | **delete length** | **Insert rate** | **insert length** | **total alignments** | **repeat alignments** | **% repeat alignments** |
| AT2E1 | 41.39844 | **90.27** | 75.28 | 292.95 | 0 | 0.29 | 0.01 | 2.34 | 0.01 | 1.92 | 90840582 | 11310095 | **12.45** |
| AT2E2 | 44.0432 | **83.67** | 69.75 | 290.88 | 0 | 0.2 | 0.01 | 2.17 | 0.01 | 1.78 | 89962866 | 12024888 | **13.36** |
| AT2E3 | 46.13089 | **81.21** | 64.78 | 288.95 | 0 | 1.08 | 0.05 | 2.82 | 0.03 | 2.42 | 94595795 | 11393092 | **12.04** |

Table S7: Summary of OMArk proteome quality assessment for longest isoforms of Warnstorfia fluitans

| Item | Count | % |
| --- | --- | --- |
| Completeness assessment |  |  |
| Ancestral clade used |  |  |
| Number of conserved HOGs |  |  |
| Complete | 5363 | 86.96 % |
| Single: | 2006 | 32.53 % |
| Duplicated: | 3357 | 54.43 % |
| Duplicated, Unexpected | 3153 | 51.13 % |
| Duplicated, Expected | 204 | 3.31 % |
| Missing: | 804 | 13.04 % |
| Consistency assessment |  |  |
| Number of proteins | 28391 |  |
| Total consistent | 24525 | 86.38 % |
| Consistent, partial hits | 4583 | 16.14 % |
| Consistent, fragmented | 615 | 2.17 % |
| Inconsistent lineage placements |  |  |
| Total inconsistent | 1209 | 4.26 % |
| Inconsistent, partial hits | 430 | 1.51 % |
| Inconsistent, fragmented | 149 | 0.52 % |
| Contaminants |  |  |
| Total contaminants | 0 | 0 % |
| Contaminants, partial hits | 0 | 0 % |
| Contaminants, fragmented | 0 | 0 % |
| Total Unknown | 2657 | 9.36 % |
| Species composition |  |  |
| Clade | *Physcomitrium patens* |  |
| No. associated query proteins | 25734 | 90.64 % |

Table S8: Coverage of 135 pairs of duplicated gene-flanking regions (provided in a separate file). Coverage of highly similar pairs (i.e., those marked in read boxes in Fig. S2) is highlighted in purple.
