## Supplementary figures and images for "*De novo* genome assembly of the floating-hook moss *Warnstorfia fluitans*"

### Fig. S2

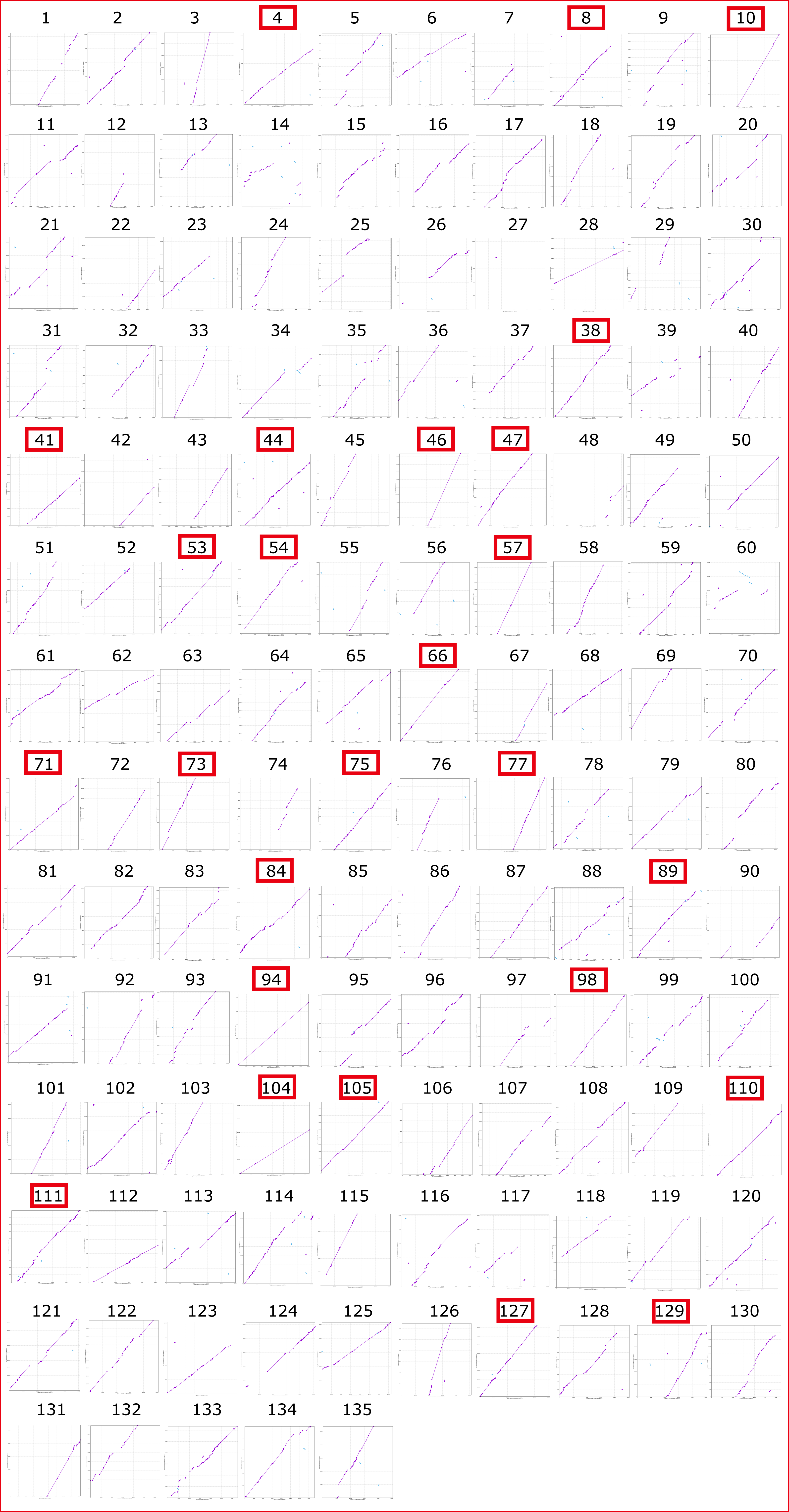
1 2 3 4 5 6 7 8 9 10
11 12 13 14 15 16 17 18 19 20
21 22 23 24 25 26 27 28 29 30
31 32 33 34 35 36 37 38 39 40
41 42 43 44 45 46 47 48 49 50
51 52 53 54 55 56 57 58 59 60
61 62 63 64 65 66 67 68 69 70
71 72 73 74 75 76 77 78 79 80
81 82 83 84 85 86 87 88 89 90
91 92 93 94 95 96 97 98 99 100
101 102 103 104 105 106 107 108 109 110
111 112 113 114 115 116 117 118 119 120
121 122 123 124 125 126 127 128 129 130
131 132 133 134 135
